# Calcium flux through ER-TGN contact sites facilitates cargo export

**DOI:** 10.1101/2022.12.19.521097

**Authors:** Bulat R. Ramazanov, Rosaria Di Martino, Abhishek Kumar, Anup Parchure, Yeongho Kim, Oliver Griesbeck, Martin A. Schwartz, Alberto Luini, Julia von Blume

## Abstract

Ca^2+^ influx into the trans-Golgi Network (TGN) promotes secretory cargo sorting by the Ca^2+^-ATPase SPCA1 and the luminal Ca^2+^ binding protein Cab45. Cab45 oligomerizes upon a local Ca^2+^ influx, and Cab45 oligomers sequester and separate soluble secretory cargo from the bulk flow of proteins in the TGN. However, how this Ca^2+^ flux into the lumen of the TGN is achieved remains elusive, as the cytosol has a very low steady-state Ca^2+^ concentration. The TGN forms membrane contact sites (MCS) with the Endoplasmic Reticulum (ER), whereby the close apposition of the two organelles allows the protein-mediated exchange of molecular species such as lipids. Here we show that the TGN export of Cab45 clients requires the integrity of ER-TGN MCS and IP3R-dependent Ca^2+^ fluxes in the MCS, suggesting Ca^2+^ transfer between these organelles. Using an MCS-targeted Ca^2+^ FRET sensor module, we measure the Ca^2+^ flow in these sites in real-time. These data show for the first time that ER-TGN MCS facilitates Ca^2+^ transfer required for SPCA1-dependent cargo sorting and export from the TGN, thus solving a fundamental question in cell biology.

**Summary:** The current study demonstrates that the trafficking of COMP and LyzC relies on Ca^2+^ flux between the endoplasmic reticulum (ER) and trans-Golgi Network (TGN). This process requires the activity of IP3 receptors, present in ER membranes, and depends on the integrity of the membrane contact site between these two organelles.

## Introduction

Protein secretion is a fundamental process and facilitates the integrity and cell-cell communication of multiple-tissue organisms (Uhlen et al., 2015). Secreted proteins are synthesized in the Endoplasmic Reticulum (ER) and transported to the Golgi apparatus in COPII coated vesicle (Barlowe and Miller, 2013; Gillon et al., 2012; Zanetti et al., 2013). Upon reaching the Golgi apparatus, these cargo molecules further transit from the *cis* to trans-Golgi cisterna and finally get the trans-Golgi Network (TGN) (De Matteis and Luini, 2008; Di Martino et al., 2019; Guo et al., 2014; Kienzle and von Blume, 2014). At the TGN, these proteins are sorted and packed into different transport carriers to reach their destination, such as the cell surface, the endosomal system, or secretory granules in specialized cells (Mostov and Cardone, 1995; Stalder and Gershlick, 2020; Tang, 2001).

Protein sorting at the TGN is a complex process involving physical features of the cargoes themselves and elements of the local environment and so remains a poorly understood process for many proteins. Understanding how soluble secretory proteins can be accurately sorted into transport vesicles targeting the plasma membrane is challenging. No cargo receptors connect these soluble molecules to the TGN membrane (Kienzle and von Blume, 2014; Pakdel and von Blume, 2018; Ramazanov et al., 2021).

Work from us and others has shown that Calcium (Ca^2+^) is a significant regulator of cargo sorting at the TGN. The Secretory Pathway ATPase 1 (SPCA1) pumps Ca^2+^ from the cytoplasm into the TGN lumen in an ATP-dependent manner (Kienzle et al., 2014; Lebreton et al., 2021; Lissandron et al., 2010; Missiaen et al., 2007; Pizzo et al., 2011; Pizzo et al., 2010; Sepulveda et al., 2008; von Blume et al., 2011; Wong et al., 2013). In response to luminal Ca^2+^ influx, the Golgi resident protein Cab45 oligomerizes and captures cargo molecules before they are packed into sphingomyelin-rich vesicles that bud from the TGN (Crevenna et al., 2016; Deng et al., 2018; Scherer et al., 1996; von Blume et al., 2012). Despite the importance of Ca^2+^ in these processes, the source of the Ca^2+^ pumped into the TGN lumen by SPCA1 is unknown, as the cytosolic Ca^2+^ concentrations are in the low nanomolar range (Berridge et al., 2003).

The TGN forms MCSs with the ER, where lipids are transferred to mediate the non-vesicular inter-organelle communication (Masone et al., 2019; Venditti et al., 2020; Venditti et al., 2019a). Endoplasmic reticulum to TGN membrane contact sites (ER-TGN MCSs) contain tethering proteins such as vesicle-associated membrane proteins A and B (VAPA and VAPB) and lipid transfer proteins such as Oxysterol-binding protein 1 (OSBP1) (Lehto and Olkkonen, 2003). OSBP1 possesses dual organelle targeting motifs as it can bind the ER proteins VAPA and VAPB through an acidic track (FFAT) domain. It is also able to attach to the TGN through the Pleckstrin homology (PH) domain, which recognizes Phosphatidylinisitol-4-phosphate (PI4P) or Arf1-GTP on TGN membrane (Kawano et al., 2006; Kumagai and Hanada, 2019; Mesmin et al., 2017). At the ER/TGN interface, OSBP1 counter transports PI4P from the TGN and cholesterol from the ER in a process mediated by the oxysterol-binding domain (OBD) of OSBP1 that binds PI4P or cholesterol in a mutually exclusive manner (Mesmin et al., 2013).

Recent studies report the importance of lipid transfer in ER-TGN MCSs in regulating protein export from the TGN (Wakana et al., 2021). In addition to their implication in lipid transfer, MCSs allow Ca^2+^ transfer between the ER and other organelles, such as the mitochondrion (Kelly, 1985; Pfeffer and Rothman, 1987; Rizzuto et al., 1993; Rizzuto et al., 1998). We, therefore, hypothesized that ER-TGN MCSs could provide Ca^2+^ to facilitate SPCA1 and Cab45-dependent cargo sorting at the TGN. Here we show that the trafficking of Cab45 clients relies on the IP3 receptor (IP3R), which is present in ER membranes, and on the integrity of the ER-TGN MCSs.

Furthermore, we generated an MCS-specific sensor to measure changes in Ca^2+^ levels within these sites, revealing that this Ca^2+^ flux plays an essential role in cargo export from the TGN and depends on tethering between ER and TGN. With these data, we solve a central unresolved question in cell biology.

## Results and discussion

### Inhibition of IP3R delays TGN export of Cab45 clients

Our previous work demonstrates that sorting soluble secretory proteins requires a transient SPCA1-mediated Ca^2+^ influx into the TGN to facilitate Cab45 oligomerization (Crevenna et al., 2016; Deng et al., 2018; von Blume et al., 2012). However, the cytosolic Ca^2+^ concentrations at steady state are in the low nanomolar range and thus may not provide the amount of Ca^2+^ ions required to promote the sorting of Cab45 clients (Crevenna et al., 2016; Pizzo et al., 2011). With the ER being the largest Ca^2+^ store within the cell, we hypothesized that the MCS between the ER and the TGN is a likely candidate to provide Ca^2+^ for sorting at TGN, which could be mediated through an IP3 receptor (IP3R)-dependent mechanism.

To investigate if the release of Ca^2+^ from the ER has an impact on the sorting and export of soluble secretory cargo molecules from the TGN, we analyzed trafficking and secretion of the well-established Cab45-clients: cartilage oligomeric protein (COMP) and Lysozyme C (LyzC) (von Blume 2011; von Blume 2012) in the presence and absence of the IP3R antagonist (2-APB) (Maruyama et al., 1997). We used the RUSH (Retention Using Selective Hooks) system to quantify the trafficking and packaging of COMP or LyzC, respectively, into secretory vesicles in HeLa cells in the presence of 2-APB (70 μM) or DMSO (control) (Boncompain et al., 2012). To this end, HeLa cell lines were transfected with RUSH constructs containing COMP-EGFP or LyzC-EGFP to analyze intracellular trafficking of EGFP-fused proteins at different time points after synchronous release from the ER by biotin addition (**Figure 1A**). We observed simultaneous export of COMP-EGFP from the ER in control and 2-APB treated cells (**Figure 1B [0, 20 min]**). However, the appearance of cytosolic vesicles (TGN carriers) was significantly delayed in cells treated with 2-APB compared to control cells at later time points (**Figure 1B, C [30, 40, 60 min]**).

**Figure 1.**
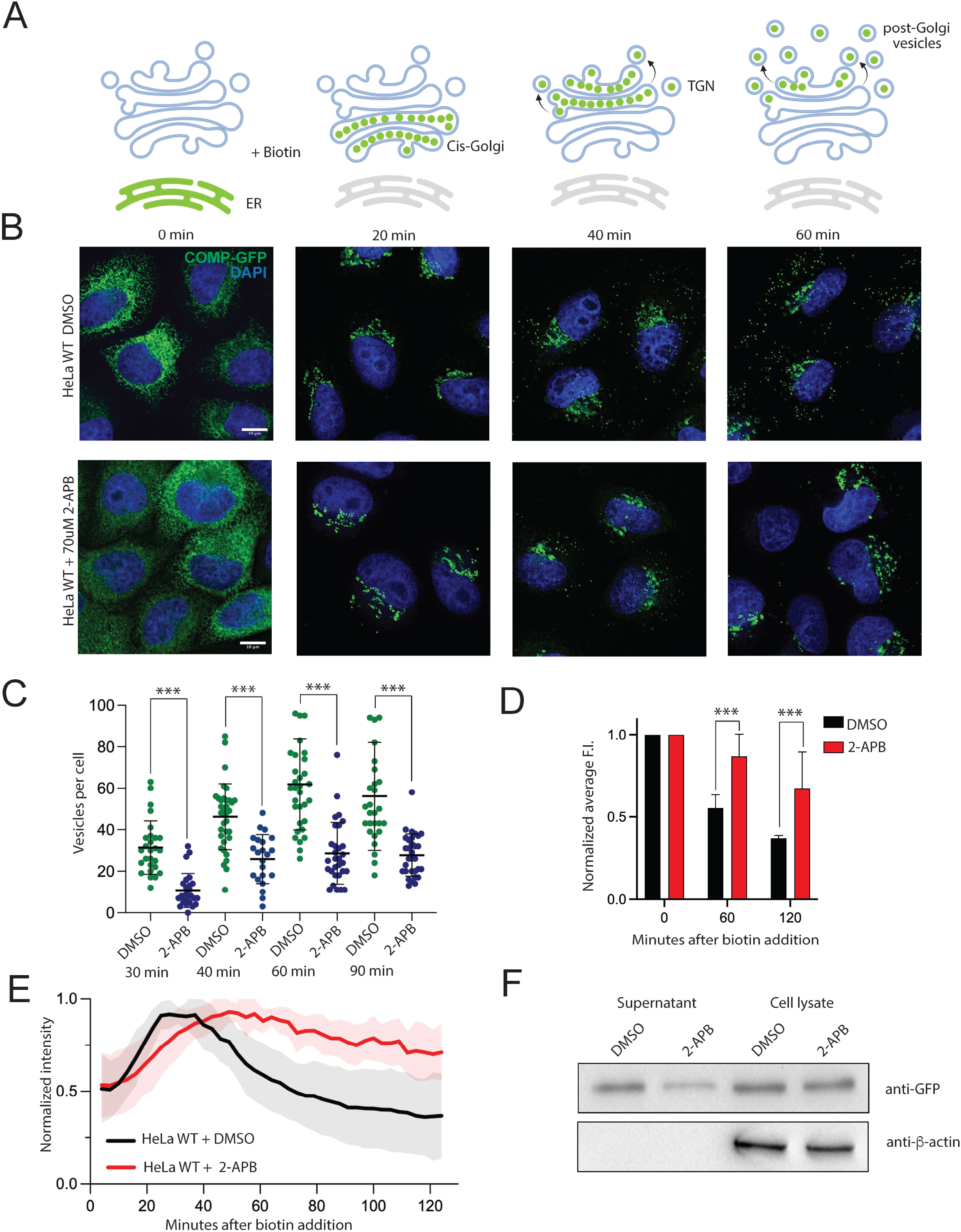
Inhibition of IP3R activity delays TGN export of Cab45 clients. (A) Schematic representation of the Retention Using Streptavidin Hooks (RUSH) assay with fluorescent tagged client molecules. (B) Representative immunofluorescence images of the RUSH experiments showing COMP-EGFP transport in HeLa lines treated with DMSO and 70 nM 2-APB. HeLa cells were transfected with KDEL-IRES-SBP-COMP-EGFP and fixed at 0, 20, 40, and 60 min after the addition of biotin. Z-stack images (d = 0.2 μm) were analyzed. The arrowheads indicate cytoplasmic vesicles. Scale bars, 10 μm. (C) The numbers of COMP budding vesicles from RUSH experiments with KDEL-IRES-SBP-COMP-EGFP in HeLa lines treated with DMSO and 2-APB were quantified. The cytoplasmic vesicles were counted at each time point by analyzing z-stack images (d = 0.2 μm). Scatter dot plot represents the means ± SD of at least three independent experiments (n > 30 cells per condition). Statistical test, Kruskal–Wallis. (D) Plot representing normalized average fluorescence intensity of COMP-EGFP in cells by FACS at 0, 60 and 120 minutes of RUSH experiments. (E) Plot representing normalized fluorescence intensity of LyzC-EGFP within TGN (ROI was defined by GALNT1 area) in cells treated with DMSO and 2-APB. (F) Western blot showing LyzC-GFP in Hela cells treated with DMSO and 2-APB in cell lysates and in secreted medium (top) and β-actin as a loading control.

To confirm the observed phenotype on a cell population, we performed RUSH experiments using COMP-EGFP as cargo in HeLa cells treated with DMSO (control) or 2-APB with subsequent FACS analysis. We used this assay to quantify the intracellular accumulation of EGFP-COMP in DMSO or 2-APB-treated cells. Cells were fixed at 0, 30, 60, and 120 min after biotin addition, and 10^4^ cells for each time point were analyzed by FACS. We calculated the average arithmetical value for the fluorescent intensity of COMP-EGFP obtained from each sample’s green emission fluorescence channel. The arithmetical average values for fluorescence intensity of COMP-EGFP from 2-APB treated cells were 1.8 and 2.1-fold higher than in control cells after 60 and 120 minutes after biotin addition, respectively, indicating a more extended residence of the EGFP-COMP inside cells in 2-APB treated cells confirming our microscopy observations (**Figure 1D, Supplementary S1A, S1C**).

To correlate these results with the actual TGN exit of the cargo molecules, we applied live-cell imaging of the exiting EGFP-tagged cargo molecules in the presence of the GALNT1-BFP TGN marker. We measured time-dependent changes in fluorescence intensity of EGFP-tagged protein within the ROI of TGN defined by the GALNT1-BFP signal. These results showed that the reduction in the number of vesicles in 2-APB-treated cells in the RUSH experiments was consistent with a prolonged residence of cargo in the TGN compared to DMSO-treated cells (**Figure 1E**). To quantify this phenotype, we applied a non-linear regression function of the intensity values on the plot shown in Figure 1D (**Supplementary S1E**). In addition, we calculated span values for each curve representing changes in LyzC-EGFP intensity within Golgi ROI. The span was defined as the difference between the fluorescence intensity of EGFP at the starting point and the predicted plateau for each curve, representing a change in fluorescence intensity during the experiment. The span value calculated for LyzC-EGFP expressing cells in the presence of 2-APB exhibited a two-fold decrease compared to control cells (span value for 2-APB and DMSO samples were 0.8 and 1.5 respectively), indicating a significant defect in TGN export of LyzC-EGFP in these cells.

To further validate that the inhibition of the IP3R reduces the secretion of the Cab45 client LyzC from the cells, we performed a secretion assay. We generated stable cell lines expressing EGFP-tagged LyzC under a constitutive promoter by lentiviral transduction of HeLa cells. Secretion assays were performed in a complete growth medium, and secreted LyzC-EGFP in the supernatant was immunoprecipitated using GFP trap agarose beads before analysis by western blotting. The western blot analysis revealed a 48% reduction in secreted LyzC-EGFP in the 2-APB-treated condition compared to DMSO-treated control cells (**Figure 1F**).

These data suggested that the activity of IP3 receptors impacts the TGN export and secretion of Cab45 clients. Previous work has shown that Golgi-localized IP3 receptors have no impact on SPCA1 dependent Ca^2+^ uptake (Wong et al., 2013). Therefore we speculated that Ca^2+^ flow between these organelles might be facilitated by ER-TGN MCS (Ramazanov et al., 2021).

### MCS integrity is essential for TGN export of Cab45 clients

VAPA and VAPB proteins are essential in maintaining ER-TGN MCS integrity by tethering ER and TGN. Deleting these proteins from cells reduces contacts between the organelles (Lev, 2010; Phillips and Voeltz, 2016; Venditti et al., 2019b). To examine the role of ER-TGN contact site in the trafficking of Cab45 clients, we performed RUSH experiments and analyzed the trafficking of COMP-EGFP in VAPA/VAPB-depleted HeLa cells. To this end, HeLa cells were transfected with non-targeting siRNA as control or siRNAs targeting VAPA and VAPB. After 24 hours, HeLa cell lines were transfected with RUSH-constructs expressing COMP-EGFP.

RUSH experiments were performed, and samples were fixed at different time points after biotin addition. Immunofluorescence images captured at 30, 40, and 60 minutes after biotin addition showed that compared to control cells, VAPA/VAPB siRNA-treated cells exhibited a significant delay in the formation of COMP-EGFP containing post-Golgi vesicles (**Figure 2A [30, 40, 60 min]**). Consistent with this result, quantification of post-Golgi vesicles at different time points from a randomly selected population of control siRNA or VAPA/VAPB siRNA-treated cells revealed a decrease in the amount of TGN-derived vesicles (**Figure 2B**).

**Figure 2.**
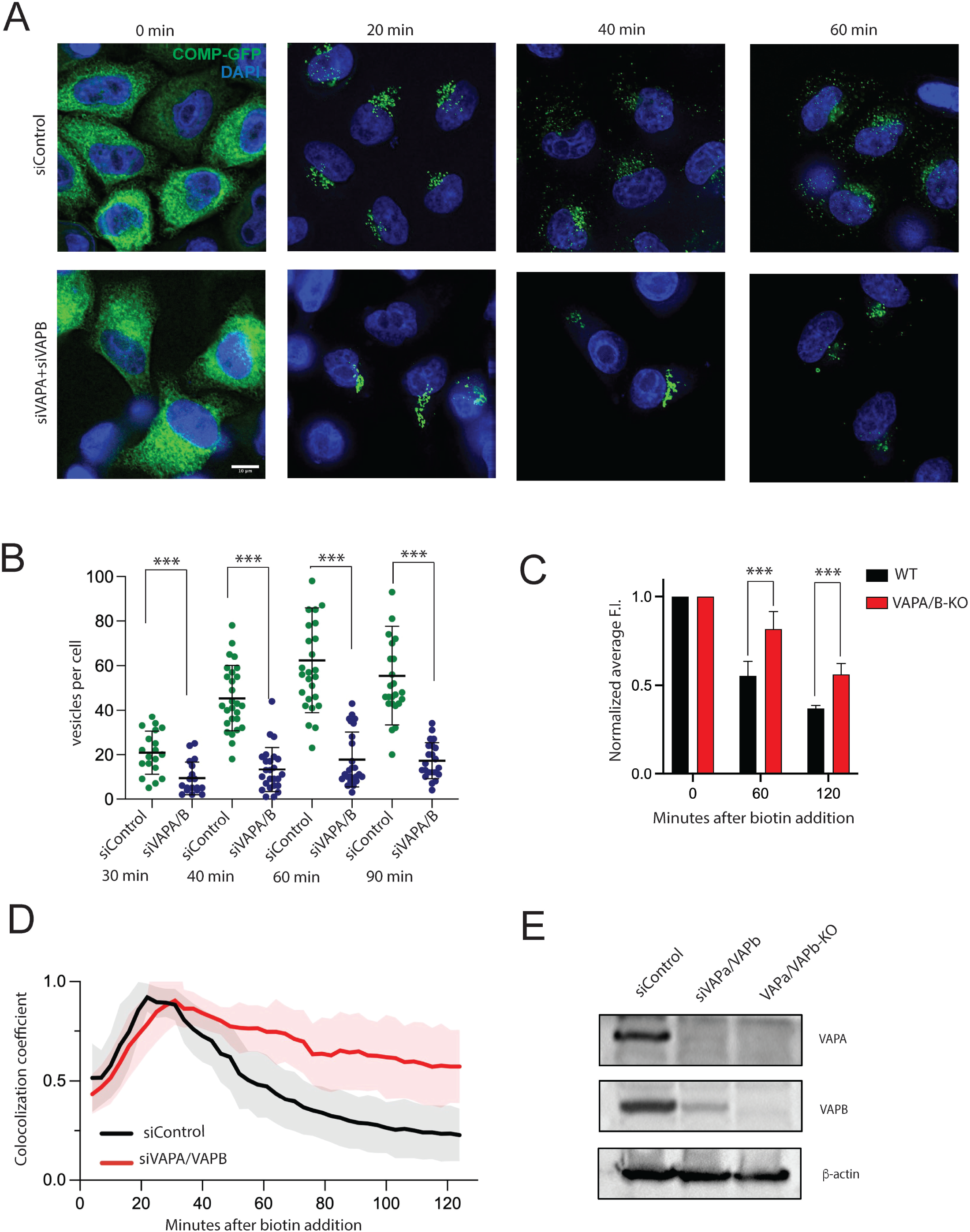
Depletion of VAPA and VAPB delays TGN export of Cab45 clients. (A) Representative immunofluorescence images of RUSH experiments showing COMP-GFP transport in HeLa lines treated with siRNA against VAPA and VAPB proteins. Cells were transfected with KDEL-IRES-SBP-COMP-EGFP and fixed at 0, 20, 40, and 60 min after the addition of biotin. Z-stack images (d = 0.2 μm) were analyzed. Scale bars, 10 μm. (B) The numbers of COMP budding vesicles were quantified. The cytoplasmic vesicles were counted at each time point by analyzing z-stack images (d = 0.2 μm). Scatter dot plot represents the means ± SD of at least three independent experiments (n > 30 cells per condition). Statistical test, Kruskal–Wallis. (C) Plot representing normalized average fluorescence intensity of COMP-EGFP in cells by FACS at 0, 60 and 120 minutes of RUSH experiments in HeLa WT and VAPA/VAPB-dKO lines (D) Plot representing normalized fluorescence intensity of LyzC-EGFP within TGN (ROI was defined by Galnt1 area) in cells treated with non-targeting (control) siRNA and siRNA targeting VAPA and VAPB. (E) Expression of VAPA and VAPB proteins in Hela WT line transfected with control (non-targeting) siRNA, Hela line transfected with siVAPA/siVAPB and Hela VAPA/VAPB-dKO line. β-actin was used as a loading control.

We confirmed the delayed export of Cab45 clients seen in VAPA and VAPB knockdown with RUSH LyzC or COMP, respectively, in control versus VAPA/VAPB-dKO HeLa cell lines (Dong et al., 2016) (see immunofluorescence **Supplementary S2A, S2B [0, 20 min]**), and quantification (**Supplementary S2C, S2D**). In parallel, we measured the time-dependent intracellular decrease of COMP-EGFP by FACS in control versus VAPA/VAPB deficient cells. The average intensity values in the GFP channel from VAPA and VAPB deficient cells were 1.5-fold higher than in control cells at 60 and 120 minutes after biotin addition, indicating accumulation of COMP-EGFP in these cells (**Figure 2C, Supplementary S1A, S1C**).

To demonstrate that the secretion defect is caused by impaired TGN export, we performed live-cell imaging of the exiting EGFP-tagged Cab45 clients in the presence of the GALNT1-BFP TGN marker. We observed time-dependent changes in LyzC-EGFP intensity within the TGN ROI and a 1.6-fold difference in span values in control versus VAPA and VAPB-depleted cells. In addition, we observed a prolonged residence of LyzC in the TGN of HeLa cell lines transfected with siRNA to VAPA and VAPB compared to control cells (**Figure 2D, Supplementary S1F**). The efficiency of knockdowns of VAPA and VAPB proteins was analyzed by western blotting (**Figure 2E**).

To investigate whether this phenotype is specific for certain soluble secretory proteins and does not affect other proteins transported by bulk flow secretion, we measured the time-dependent intracellular decrease of COMP-EGFP and EQ-sol-GFP by FACS in control versus VAPA/VAPB deficient HeLa cells as well as 2-APB treated cells. These data indicated that the secretion of EQ-sol-GFP, a non-specific marker of bulk flow secretion (Deng et al., 2016), is unaffected by VAPA and VAPB-depletion (**Suppl. S1A-C**).

Together these data showed that the TGN export of the Cab45 clients COMP and LyzC from TGN requires intact MCSs between ER and TGN (**Figure 2**) and IP3R-dependent Ca^2+^ release from the ER (**Figure 1**). These data also suggest that ER-TGN MCS could serve as potential sites for Ca^2+^ transfer.

### Targeting Twitch-based FRET sensors to ER-TGN MCS

Our next goal was to test potential Ca2+ flow at ER-TGN MCSs directly. These MCSs would be potential structures that could serve as hotspots for Ca^2+^ transfer between the organelles. To evaluate possible Ca^2+^ flows in these sites, we used Förster resonance energy transfer (FRET)-based Ca^2+^ biosensors called Twitch. These sensors contain a minimal Ca^2+^ binding moiety derived from the C-terminal domain of troponin C incorporated between mCerulean3 and cpVenuscd (Thestrup et al., 2014). To target the sensors to ER-TGN MCS in living cells, we introduced an amino acid sequence coding the N-terminal region of OSBP1, including a PH domain and an FFAT motif (**Figure 3A**). As the N-terminal disordered domain of OSBP seems to be crucial for active lipid transport in the ER-TGN MCS (Jamecna et al., 2019), we generated sensors containing a disordered domain (N-PH-FFAT) as well as a sensor without that domain (PH-FFAT) (**Supplementary S3A, S3B**). We constructed stable HeLa cell lines expressing the MCS-Twitch sensors under a doxycycline-inducible promoter to obtain stable and equal protein expression levels. We confirmed the expression of the respective sensor after doxycycline induction by immunofluorescence and live-cell imaging (**Supplementary S3B**). To investigate the correct localization of the sensors at the TGN, cells were fixed and stained with antibodies recognizing GM130 or TGN46. Immunofluorescence microscopy data confirmed that the MCS-Twitch sensors expressed in HeLa cells correctly localize (**Supplementary S3C, S3D**).

**Figure 3.**
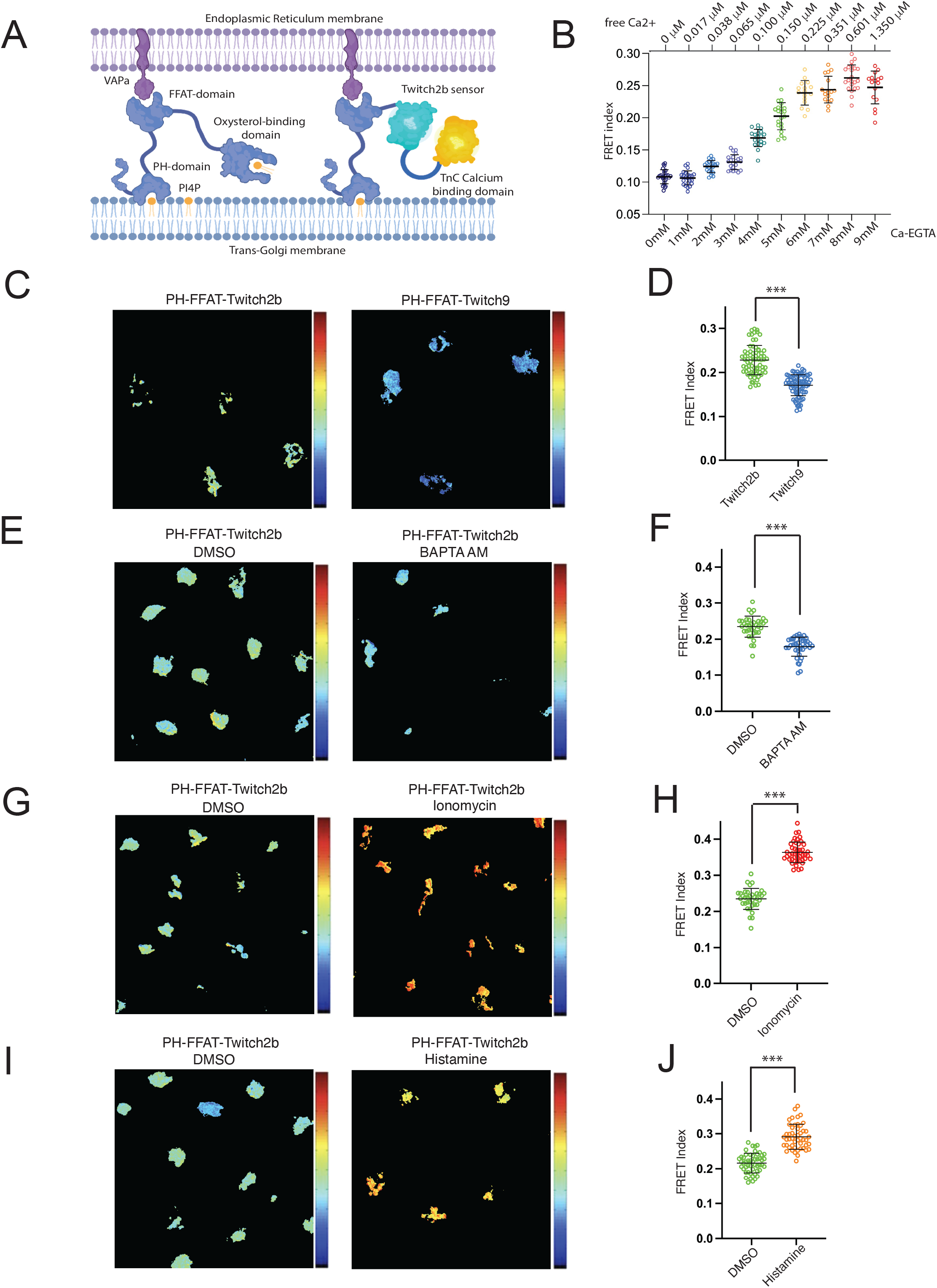
MCS targeting FRET sensor responses to changes in Ca^2+^ concentration at ER-TGN MCS. (A) Schematic representation of OSBP and Twitch2b Calcium Sensor targeting Endoplasmic Reticulum-Golgi membrane contact sites. The PH domain of OSBP binds to PI4P on the outer leaflet of the TGN membrane, and the FFAT domain interacts with VAPA on the ER membrane. (B) The calibration curve for the Twitch2b sensor used a buffer with increasing concentration of free Ca^2+^ ions, demonstrating a correlation between FRET indices and free Ca^2+^. (See Materials and Methods for a detailed description). (C) Pseudocolor map of FRET index and dot plot (D) representing FRET indices for PH-FFAT-Twitch2b and PH-FFAT-Twitch9x within Golgi ROI of live cells and at steady state (***p<0.05). (E) Pseudocolor map of FRET index and dot plot (F) representing FRET indices for PH-FFAT-Twitch2b at steady state and treated with 25uM BAPTA-AM for 20 min (***p<0.05). (G) Pseudocolor map of FRET index and dot plot (H) representing FRET indices for PH-FFAT-Twitch2b at steady state and treated with 1uM ionomycin for 20 min (***p<0.05). (I) Pseudocolor map of FRET index and dot plot (J) representing FRET indices for PH-FFAT-Twitch2b at steady state and treated with 1uM Histamine for 2 min (***p<0.05). Pseudocolor bar FRET index value in range 0-0.4.

### The Twitch2b-MCS FRET sensor detects Ca^2+^flows

To determine the range of Ca^2+^ signals detectable at the MCS at steady-state (non-treated cells incubated at 37°C), we constructed four MCS-Twitch sensors with different Ca^2+^ affinities (depicted in **Supplementary Tables 1, 2**). For quantitative FRET measurements of Ca^2+^, we calculated the FRET index value in the cell line expressing the respective sensor as an approximation of the FRET/molecule (Grashoff et al., 2010; Kumar et al., 2016). The normalized FRET index values were acquired by measuring FRET intensity, subtracting the background noise and the bleed-through for the two fluorophores, and normalizing them to FRET acceptor intensity. Measuring FRET indexes from the MCS-Twitch sensors at steady revealed that compared to MCS-Twitch9x, MCS-Twitch7x, and MCS-Twitch8x, the Twitch2b sensor showed the highest FRET index (**Supplementary S3E**).

We performed a series of control experiments to test if the Twitch2b sensor was responsive to Ca2+ perturbations in the cells. First, we determined the FRET values in HeLa cells expressing the respective MCS-Twitch sensor by live-cell fluorescence microscopy in cells treated with DMSO (control) or the Ca^2+^ chelating agent BAPTA-AM. As expected, chelating intracellular Ca^2+^ ions with BAPTA-AM significantly decreased the FRET index of MCS-Twitch2b (**Figure 3E, F**). However, the FRET indices of MCS-Twitch2b upon treating the cells with BAPTA were similar to the baseline levels of the low Ca^2+^ affinity MCS-Twitch7x, 8x and 9x sensors (**Figure C, D; Supplementary S3E**). Therefore, Twitch2b was selected for all further experiments.

To quantify the range of free Ca^2+^ concentrations, we calibrated the Twitch2b sensor. To calibrate Twitch2b in live cells, we used HeLa cells stably expressing CYTO-Twitch2b. We applied a reciprocal dilution of buffers containing increasing ratios of Ca-EGTA/K2-EGTA concentrations, and the free Ca^2+^ ion concentrations were calculated as described in Materials and Methods (**Supplementary S4A, S4B**). The FRET indexes obtained from Twitch2b at different ratios of Ca-EGTA/K2-EGTA in the calibration experiment allowed us to build a calibration curve that demonstrated a strong correlation between the FRET index value and the concentration of free Ca^2+^ (calibration plot shown in **Figure 3B**). Thus, we developed a powerful tool to measure Ca^2+^ levels in the ER-TGN-MCS. We developed a tool that accurately measures Ca2+ levels in the ER-TGN-MCS.

To measure the effects of increased intracellular Ca^2+^ concentrations on the FRET signals from MCS-Twitch2b sensors, we utilized active and passive means of increasing cytosolic Ca^2+^ levels. Treatment of cells with ionomycin, which raises the intracellular Ca^2+^ level, significantly increased the FRET index (Figure 3G, H). Several signaling pathways, including cell surface receptors, are known to utilize Ca^2+^ ions as second messengers for the downstream signaling (Carafoli, 2002; Dickenson and Hill, 1994; Thillaiappan et al., 2017). Activation of Histamine receptors (H1-receptor) at the plasma membrane causes the activation of PLC, which in turn elevates intracellular Ca^2+^ through an IP3-dependent mechanism. To test if the MCS sensor detects these signals, we performed the FRET measurements in HeLa cells expressing PH-FFAT-Twitch2b incubated with either DMSO (control) or after treatment with histamine. (**Figure 3I, J**). The data showed an increase in the FRET index– revealing a physiological link between MCS Ca^2+^ levels and the signaling receptor IP3.

Because we did not observe differences in the FRET index between PH-FFAT-Twitch and N-PH-FFAT-Twitch sensors (**Supplementary 3E, 3F, 3G, 3H**), we decided to perform further experiments using the PH-FFAT targeting motif.

The data indicates that the release of Ca^2+^ caused by IP3R stimulation leads to an increase in Ca^2+^ levels in MCS and that our sensor sensitively detects these changes.

### TGN protein abundance and Ca^2+^flux at MCS are coupled

Our previous work showed that TGN Ca^2+^ influx is necessary for the TGN exit of secretory proteins (Crevenna et al., 2016; Deng et al., 2018; Kienzle et al., 2014; von Blume et al., 2011). Therefore, we hypothesized that there must be a correlation between cargo influx into the TGN and Ca^2+^ flow in the MCS. To test if Ca^2+^ in ER-TGN MCSs is influenced by protein abundance, we treated HeLa cells expressing MCS-Twitch 2b with Cycloheximide (CHX) to block *de novo* protein synthesis. We incubated cells for 1, 2, and 4 hours and analyzed FRET signals. We observed a time-dependent decrease in the FRET index of cells treated with CHX. To further demonstrate that this is correlated with cargo abundance in the TGN, we incubated HeLa cells expressing MCS-Twitch 2b at 20°C to arrest secretory proteins in the TGN (Ladinsky et al., 2002; Matlin and Simons, 1983). Notably, FRET values were significantly decreased in cells incubated at 20°C (**Figure 4A, B**). However, duration of the 20°C block for more than 1 hour did not affect the average FRET index values (**Figure 4C**). More importantly, incubation of cells at 37°C after the 20°C-block resulted in a complete recovery of the FRET index (**Figure 4B, C)**. To validate that the changes in the FRET indices were not due to other factors, cells were incubated at 37°C and 20°C in the presence of ionomycin (**Figure 4E**). Independent of the incubation temperature, ionomycin treatment led to a recovery of FRET indexes, demonstrating that the effects are specific to Ca^2+^. These data supported that the presence of newly synthesized protein stimulates the Ca2+ flux in the MCS. Furthermore, we showed that this is specific to the cargo abundance in the TGN.

**Figure 4.**
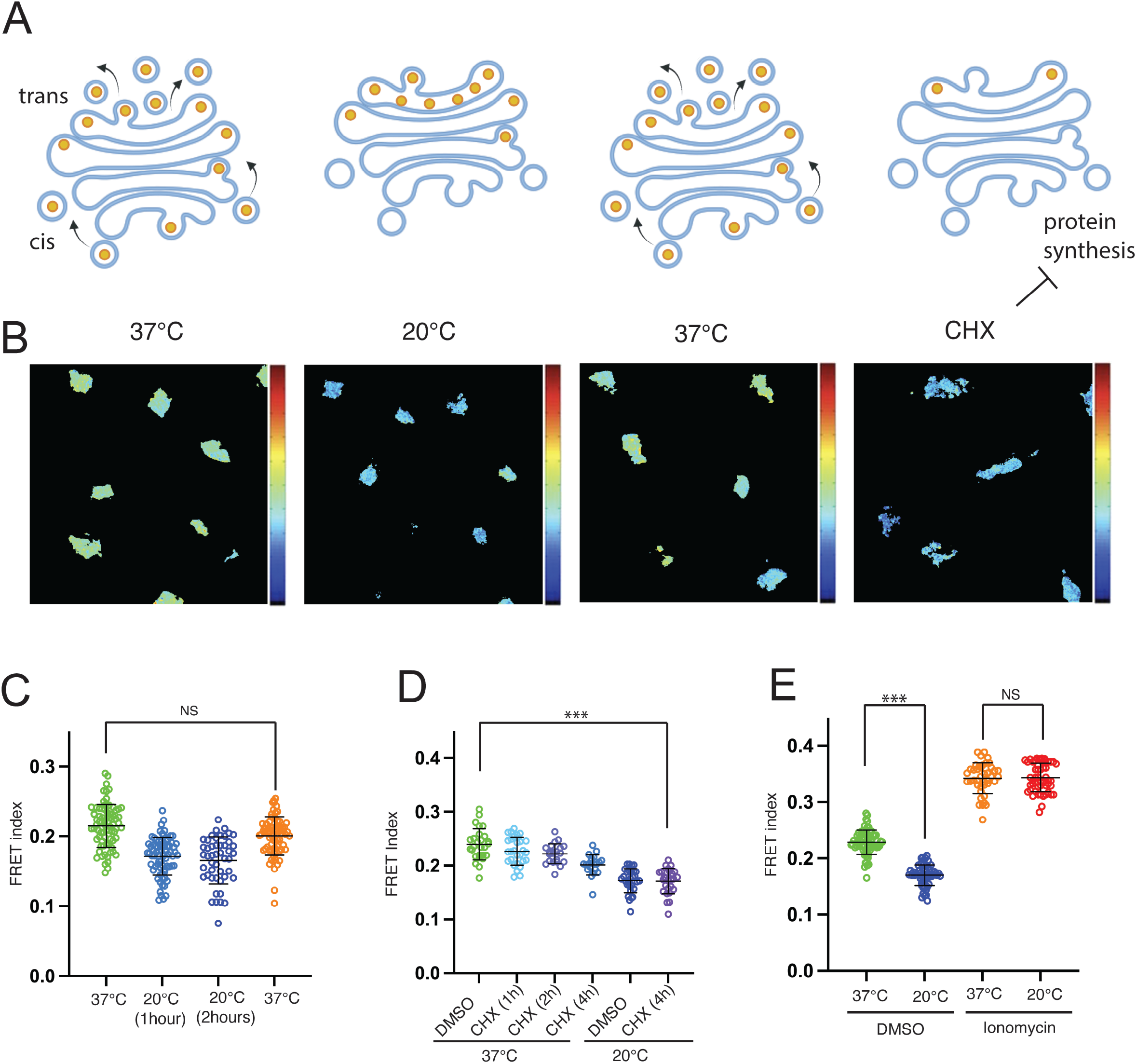
Ca^2+^ flux at ER-TGN MCS is coupled to protein trafficking. (A) Schematic representation of experiment and effect on protein trafficking in live cells during 20°C block and cycloheximide (CHX) treatment. (B) Pseudocolor maps of FRET index for PH-FFAT-Twitch2b within Golgi ROI of live cells at described above conditions. Pseudocolor bar FRET index value in range 0-0.4. (C) Dot plot representing FRET index for PH-FFAT-Twitch2b at steady state, after 20°C block for 1 and 2 hours and after 10 min recovery at 37°C (***p<0.05). (D) Dot plot representing FRET index for PH-FFAT-Twitch2b at steady state, treated with CHX for 1, 2, 4 hours and after 1 h 20°C block. (***p<0.05), (E) Dot plot representing normalized FRET index for PH-FFAT-Twitch2b at steady state, after 1 h 20°C block and effect of ionomycin at 37°C and 20°C conditions (***p<0.05).

In the current study, we quantify the abundance of Ca^2+^ flux in the ER-TGN MCS, which has remained unknown. We also demonstrate that cargo entering the TGN elicits a Ca^2+^ release in the MCS required for sorting Cab45 clients into a TGN-derived carrier (**Figure 5A**).

**Figure 5.**
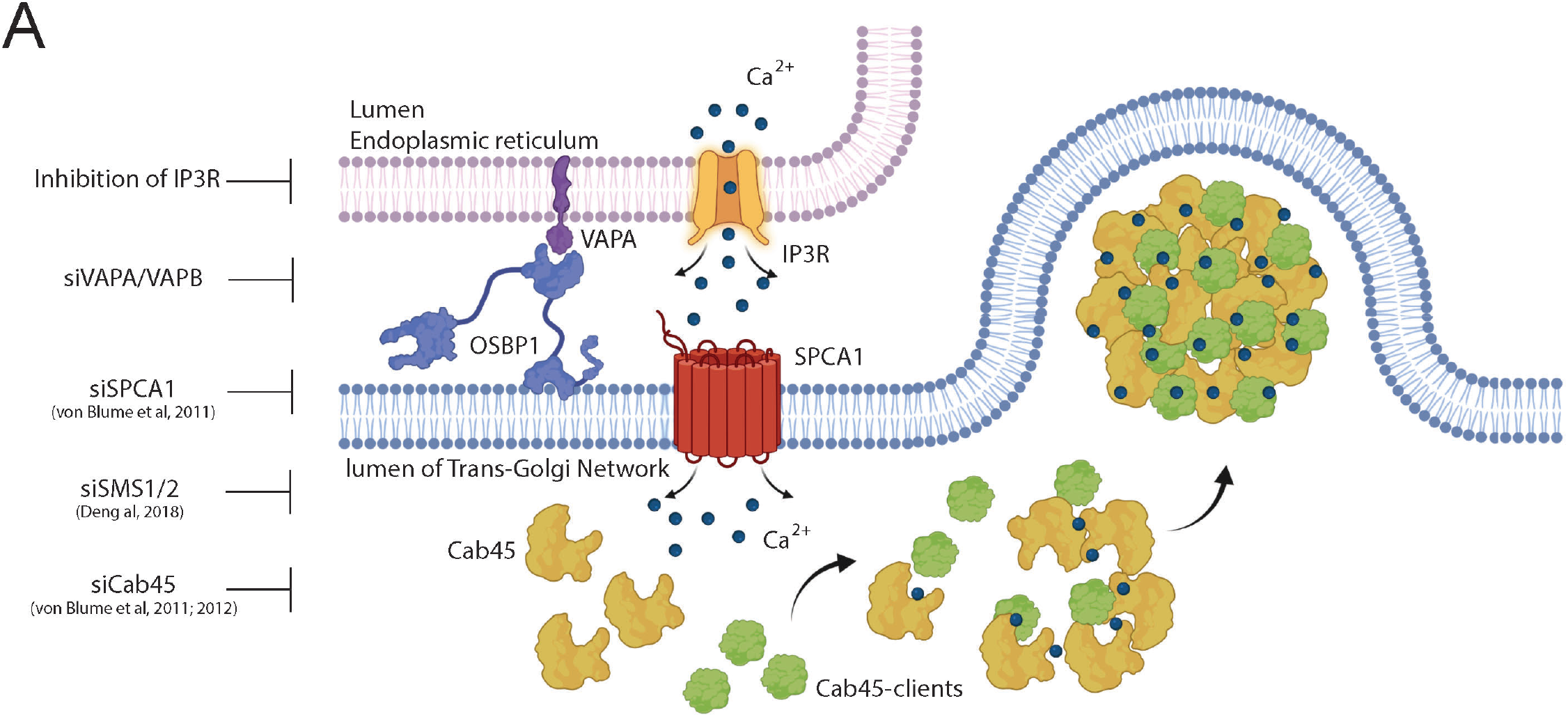
The model depicting the role of contact sites and IP3R-dependent release of Calcium ions for SPCA1-dependent sorting at TGN.

Our previous work showed that SPCA1, the only Ca^2+^ ATPase in the TGN, is required for secretory cargo sorting in the TGN (von Blume et al., 2011). Interestingly, these studies also exemplified that the Ca^2+^ binding protein Cab45 oligomerizes and forms a scaffold for collecting its clients, including LyzC and COMP (Crevenna et al., 2016; von Blume et al., 2012). Furthermore, sphingomyelin produced by SMS1 promotes SPCA activity and Cab45/client export from the TGN (Deng et al., 2018). However, the source of Ca^2+^ that drives the process has remained unknown (Ramazanov et al., 2021). In the current work, we demonstrate that the TGN export of COMP and LyzC requires the integrity of the ER-TGN MCS and IP3R-dependent Ca^2+^ fluxes in the MCS, suggesting that Ca^2+^ transfer between these organelles is the primary source for SPCA1.

The autoregulation of the secretory pathway by cellular signaling remains mysterious. However, central principles have been uncovered in recent years (Di Martino et al., 2019). On the other hand, the signaling pathway that stimulates the Ca^2+^ release through the contact site remains unknown. Our work shows that it is directly linked to IP3 production and activation of IP3R close to ER-TGN contact sites. Although current work provides an understanding of upstream signaling events that regulate of sorting of soluble proteins at TGN and depend on the activity of SPCA1, the possibility that LyzC and COMP themselves could serve as a stimulus for the activation of such pathways remains to be investigated.

## Materials and Methods

### DNA techniques and plasmid construction

Restriction enzymes for molecular biology were obtained from New England Biolabs. PCRs were performed with a Phusion Polymerase (Thermofisher) and a Mastercycler Nexus (Eppendorf). All plasmids used in this study bear ampicillin resistance for selection in *Escherichia coli* and are listed in Table 1, where the transgenes and inserts are described. The DNA sequences encoding PH-FFAT/N-PH-FFAT domains of OSBP1 fused with Twitch sensors were integrated into the donor plasmid of the transposon-based piggyBac system for stable transgene-expressing cell lines generation (Li et al., 2013). The piggyBac backbone vector (PB-T-PAF) and PB-RN and PBase were a gift from James Rini, University of Toronto, Ontario, Canada (Li et al., 2013). In brief, to generate PB-T-PAF-PHFFAT-Twitch constructs, the PB-T-PAF vector was linearized with NheI and NotI-HF restriction enzymes (NEB). PH-FFAT/N-PH-FFAT sequences (5’-NheI/3’-AscI) were amplified by PCR from pLJM1-FLAG-GFP-OSBP plasmid (Addgene#134659). The sequences encoding Twitch (Twitch2b/Twitch7x/Twitch8x/Twitch9x) Calcium sensors were obtained from plasmids that have been generously provided by Oliver Griesbeck Lab and were amplified by adding corresponding restriction sites (5’-AscI/3’-NotI). All fragments were ligated in the PB-T-PAF backbone. All cloning experiments were conducted using Phusion High-Fidelity Polymerase and T4 ligase (Thermo Fisher Scientific) according to the manufacturer’s instructions. Similar strategies generated VAPA-Twitch2b and CYTO-Twitch2b constructs. The sequences encoding Twitch2b for these constructs were amplified by adding corresponding restriction sites (5’-NheI/3’-AscI). The VAPA fragment was amplified from plasmid coding full-length VAPA protein and was gifted from Pietro De Camilli with addition of AscI and NotI restriction sites. CYTO-Twitch2b construct was generated from VAPA-Twitch2b by replacing the VAPA sequence with short stop-codon containing fragment, annealed by using the following sequences (5’CGCGCCAGAGGAGTTTTAAGC3’ and 5’GGCCGCTTAAAACTCCTCTGG3’). pLenti-LyzC-EGFP for secretion assay lines was generated using gateway cloning reaction by amplifying LyzC-EGFP from plpcx-LyzC-EGFP and cloning into pDONR221 using BP cloning reaction and then subsequently using LR cloning reaction into the destination vector to generate the desired construct. The correct sequence of all constructs was confirmed by DNA sequencing using the SmartSeq Kit from Eurofins Genomics or KECK sequencing (Yale University).

### Cell culture and generation of stable cell lines expressing Calcium sensor constructs

Cell lines were maintained in DMEM media (Gibco) containing 10% FBS (Sigma 12306C-500ML) at 37°C and 5% CO_2_. For transfection, cells were plated in antibiotic-free media 24 hours before the procedure. DNA transfections were performed using Lipofectamine 2000 reagent according to the manufacturer’s protocol. After 8 hours, the media was replaced. Transgene expression was estimated at 48 hours after transfection. To generate cell lines stably expressing the mentioned transgenes HeLa lines were used. In brief, HeLa cells at 70% confluency were transfected with PB-T-PAF (with the corresponding transgene), PB-RN, and PBase (total DNA 1.5 μg; at ratio 8:1:1) using Lipofectamine 2000 in OptiMEM-I media. Cells were selected for 48 hours with 2 μg/ml puromycin dihydrochloride (Sigma-Aldrich) and for 7 days with 400 μg/ml G418 disulfate salt (A1720-5G, Sigma-Aldrich). To induce transgene expression, cells were incubated in the presence of doxycycline (J63805 Alfa Aesar, USA) (1ug/mL) for 24 hours. The generated lines were sorted on BD FACS Aria to exclude resistant cells without transgene expression.

HeLa cell lines stably expressing LyzC-EGFP were generated using lentiviral transduction containing pLenti-LyzC-EGFP construct with followed Blasticidin 8 ug/mL (InvivoGen) selection for 48 hours. siRNA transfections were performed using Lipofectamine RNAiMAX according to standard protocol. VAPA/VAPB-dKO lines were generated by Pietro De Camilli Lab and published previously (Dong et al., 2016).

### RUSH cargo sorting assay and fluorescent microscopy

RUSH assays were performed as described previously (Boncompain et al., 2012; Deng et al., 2018). Studied cell lines were cultured in 6-well plates (Cat#353046, Corning) on glass slides (Cat. #72290-04, EMS) and transfected with RUSH-COMP-EGFP and RUSH-LyzC-EGFP constructs using Lipofectamine 2000 according to standard protocol. At 24 hours post-transfection, cells were incubated with 40 μM d-Biotin (Sigma) ingrown media for different time points (20, 30, 40, 60, and 90 min or without d-Biotin (control). For IF slides, cells were washed once with PBS, fixed in 4% PFA (Electron Microscopy Sciences) in PBS for 10 min, and mounted on 12mm coverslips (Electron Microscopy Sciences) using ProLong Gold (Thermo Fisher Scientific). Nuclear chromatin was stained by short incubation in 2.5uM DAPI (Biolegend) solution. Acquisition of EGFP was performed using a Delta Vision system by imaging z-stacks with a step size of 0.2 μm.

We empirically measured the sizes of objects between 4 and 20 pixels for quantification of vesicles using the Analyze Particles function in ImageJ, which detects vesicular carriers but omits larger objects such as the Golgi. While small-fragmented and isolated Golgi structures could be detected in error. Furthermore, only vesicles of cells expressing the RUSH construct were counted. The Fiji macro count_fixed_vesicles_V1.3 (M. Pakdel), including the Particle Analyzer plug-in by Fiji, was used to determine the number of vesicles (Deng et al., 2018). Kruskal–Wallis one-way analysis of variance was used for comparisons in RUSH experiments.

Cells were cultured in six wells on glass slides for immunostaining and fixed for 10 min with 4% paraformaldehyde. After washing with PBS, cells were permeabilized for 5 min in 0.2% Triton-X 100 and 0.5% SDS in 4% BSA. After washing with PBS, cells permeabilized with Triton-X 100 were blocked with 4% BSA for 1 h. Next, cells were incubated with primary followed by the corresponding secondary antibody for 1 h at room temperature in a blocking buffer in the dark. Slides were washed three times with PBS after incubation with antibody. Glass slides were mounted with ProLong Gold (Thermo Fisher Scientific). Antibodies to TGN46 (AHP500G, Biorad) and GM130 (610822, BD) were used at dilution 1:200.

For FACS analysis cells were fixed with 4% PFA for 10 min. After fixation washed with ice-cold DPBS and dissociated using trypsin-EDTA solution. Cells were washed with DPRS three times by centrifugation at 200g for 5 min each. FACS analysis was performed using BD LSRII machine. 10000 cells were analyzed for each sample. FACS data were analyzed using FlowJo 10.5.3 for Windows.

### RUSH protein trafficking analysis

HeLa cells were seeded on Mattek dishes (p35gc-1.5-14-c, Mattek) and cultured at 37°C with 5% CO_2_. The next day or the day before imaging, cells were transfected by Lipofectamine 2000 (Invitrogen) with two plasmids expressing GalT-BFP (a trans-Golgi maker from James Rothman lab) and RUSH COMP or RUSH Lysozyme C (see above for details of the plasmids). After five to eight hours, cells were washed and incubated with DMEM with 10% FBS overnight. Before imaging, cells were washed and briefly incubated with 37°C-warmed DMEM supplemented with 10% FBS and HEPES buffer (Gibco, 21063029). Cells were then staged on the microscope as described below.

Live-cell imaging was performed using a spinning disk confocal microscope CSUXfw-06p-01 (Yokogawa) on a Nikon eclipse Ti2 (LWD NA=0.52) microscope stand with a motorized stage with stage top Piezo. sCMOS camera Photometrics Prim 95B and CFI Plan Apo Lambda 60x oil objective were used. Also, the Oko Lab temperature control system was set to 37°C, and the fluorescence (405 nm and 488 nm) was induced using an Agilent laser combiner. Images were acquired using Nikon Elements. Fluorescence images were taken every three minutes after adding biotin (see above for the methods for RUSH experiments). At each time point, the Nikon Elements stitched 6×5 fields to generate relatively large field images that could accommodate at least 10 distinguished secretion events through 90 or 120 minutes.

The time-course images generated above were imported to Fiji (ImageJ). Images containing individual cells with distinguishable Golgi and RUSH signals were cropped and stored separately. The Golgi masks were generated using GalT-BFP signals (405 nm excitation) and ImageJ Auto-threshold and imported as ROIs. The Golgi ROIs were used to measure mean RUSH intensity in the Golgi at each time point. The mean RUSH intensities were subtracted from the background signals. Then, the maximum mean value of RUSH during the time course of the single-cell images was used to divide the RUSH intensities, thus normalizing the maximal RUSH intensities in the Golgi marker set to 1. The time course images for every cell were analyzed separately and combined to generate the mean RUSH intensity in each time point and its standard deviation, as described in the Figures. Three independently cultured cells were analyzed, and their images were combined to generate the final data.

### Secretion assay of LyzC-EGFP

For the secretion assay, 5×10^5^ cells stably expressing LyzC-EGFP were plated in a 6-well plate. After 24 hours, cells were pretreated with 70μM 2-APB and DMSO. After that incubated in a grown medium containing 2-APB and DMSO for 1 hour. Cells and media were collected separately. Cells were lysed using RIPA buffer. Collected growth media was incubated overnight with GFP-trap beads at 4°C. The following day the beads were washed 4 times with DPBS, and then protein was eluted from the beads by boiling it in 2X Laemmli SDS sample buffer (Biorad). Cell lysates and IP fractions were analyzed by western blotting.

### Live cell imaging and FRET analysis

For live cell imaging, DMEM without phenol red containing 4.5 mg ml – glucose, 25 mM HEPES, and 2 mM glutamine (Life Technologies) supplemented with 10 % FBS was used. Cells were seeded on Glass bottom dishes (D35-14-1.5-N, Cellvis) at a density 5×10^4^ per dish. The next day to induce transgene expression, doxycycline at a final concentration of 1ug/mL was added. After 24 hours, doxycycline was removed, and cells were incubated in imaging media for an additional 24 hours. ImageJ (National Institutes of Health) was used for basic image processing. All analyses were done using custom-written software (MATLAB R2014a; MathWorks). To manipulate intracellular Ca^2+^ levels as well as induce Ca^2+^ flux following drugs at corresponding concentrations were used: calcium ionomycin (I3909-1ML, Sigma) at 1μM; 2-APB (100065-100MG, Millipore) at 70μM, Histamine (H7125-1G, Sigma) 1 μM solution, BAPTA-AM (126150-97-8, Millipore) at 25 μM final concentration. Cells were treated with cycloheximide to inhibit protein synthesis at a concentration of 10ug/mL (DSC81040-5, Dot Scientific Inc).

### FRET sensor calibration experiment

Free Ca^2+^ calibration solutions were made using Calcium Calibration Buffer Kit #1 (Cat. No. C3008MP, Biotium) according to standard protocol. To perform the FRET calibration experiment, the HeLa cell line stably expressing CYTO-Twitch2b was used. 1×10^5^ cells were plated into Glass bottom dishes (D35-14-1.5-N, Cellvis), and doxycycline was added to the media at a final concentration of 1 ug/mL. After 24 hours, each plate was washed twice with 10mM EGTA, 100mM KCl, and 10mM MOPS (pH 7.2). To chelate residual Ca^2+^, cells were incubated in 10mM EGTA, 100mM KCl, and 10mM MOPS (pH 7.2) with 1 μM ionomycin for 20 min at RT. 2ml of stock solutions with free Ca^2+^ concentrations ranging from 0 μM to 39 μ μM were added to cells, and FRET indexes were measured for each condition

### FRET imaging and analysis

These analyses were done as previously described by (Kumar et al., 2016). High-resolution live FRET imaging was performed on Nikon Eclipse Ti widefield microscope equipped with a cooled charged-coupled device Cool SNAP HQ2 camera, using a ×100, 1.49 NA oil objective at 37 °C. Images were acquired using Micromanager software. Three sequential images with 500 ms exposure time were acquired with the following filter combinations: donor (Teal) channel with 460/20 (excitation filter-ex), T455lp (dichroic mirror-di), and 500/22 (emission filter-em); FRET channel with 460/20 (ex), T455lp (di) and 535/30 (em); and acceptor (Venus) channel with 492/18 (ex), T515lp (di) and 535/30 (em) filter combinations. All filters and dichroic were purchased from Chroma Technology. For data analysis, donor leakage was determined from HeLa cells transiently transfected with Vinculin-Teal, whereas acceptor cross excitation was obtained from Vinculin-Venus transfected cells. For all the calculations, respective background subtraction, illumination gradient, and pixel shift correction were performed, followed by three-point smoothening. The slope of the pixel-wise donor or acceptor channel intensity versus FRET channel intensity gives leakage (x) or cross-excitation (y) fractions, respectively. FRET map and pixel-wise FRET index for the sensors were determined from FRETindex = [FRETchannel-x(Donorchannel)-y(Acceptorchannel)]/[Acceptorchannel] ImageJ (National Institutes of Health) was used for basic image thresholding. Mean FRET index per cell was calculated for each region within the mask. Then, a student t-test was performed between the two groups to calculate statistical significance and p-value. At least P < 0.05 was considered significant. All analyses were done using custom-written software (MATLABR2020b; MathWorks) (Kumar et al., 2016).

### siRNA delivery and western blotting

Knockdown of VAPA and VAPB proteins was performed using siRNA (VAPA: 5’-AACTAATGGAAGAGTGTAAAA-3’; VAPB: 5’-AAGAAGGTTATGGAAGAATGT-3’)(Wakana et al., 2021). Non-targeting siRNA was purchased from Qiagen (Catalog No. – 1027281) and used as a negative reference control. The knockdown of VAPA and VAPB proteins was achieved by combined transfection (siVAPA and siVAPB) using LipofectRNAiMAX according to standard protocol. Knockdown efficiency was confirmed by Western blotting of cell lysates in radioimmunoprecipitation assay (RIPA) buffer (50 mM Tris-HCl, pH 7.4 [American Bioanalytical], 150 mM NaCl [American Bioanalytical], 1% Triton X100 [American Bioanalytical], 1% sodium deoxycholate [Sigma-Aldrich], and 0.1% SDS [American Bioanalytical] in Milli-Q water); protease and phosphatase inhibitor (Thermo Fisher Scientific) was added just before extraction. The cell lysate was resolved using SDS-PAGE and transferred to the nitrocellulose membrane (Biorad Laboratories) using a transfer system (Trans-Blot Turbo; Bio-Rad Laboratories). The membrane was blocked using 5% skimmed milk (American Bioanalytical) in TBS with 0.1% Tween 20 (TBST) for 1 hour and incubated with the following primary antibodies diluted in 5% milk with TBST overnight at 4°C: Anti-β-actin (1:5000, A5441-.2ML Sigma); Anti-VAPA (1:1000, SAB1402460-100G, Sigma); Anti-hVAPB (1:2000, MAB58551 R&D systems); anti-GFP (1:1000, 11814460001, Roche). The membrane was washed 3 times with 5% milk TBST and incubated with HRP-conjugated secondary antibodies (1:5000, 32230/32260; Invitrogen). Data were visualized using chemiluminescence detection on ChemiDoc Touch (Bio-Rad Laboratories).

### Graphical data and image design

Graphs were plotted in GraphPad Prism version 9.2.0 for Mac, GraphPad Software, San Diego, California, USA. Images were compiled using Adobe Illustrator 2022 (Adobe Inc. (2022). Adobe Illustrator. Retrieved from https://adobe.com/products/illustrator). Schemes were designed using Biorender software (biorender.com).

## Acknowledgments

We thank Dr. Pietro de Camilli for providing VAPA/VAPB-dKO cell lines and the Burd lab for discussions and support. Julia von Blume is funded by a Yale-start grant and by the National Institute of General Medical Sciences of the NIH under the award number GM134083-01, an Administrative Supplement 3R01GM134083-03S1 and a Project and Feasibility award from Yale Diabetic Research Center (GR112420).

## Author Contributions

B. R. Ramazanov and J. von Blume designed the experiments. B. R. Ramazanov carried out the experiments and analyzed the data. A. Kumar and M. Schwartz supported FRET data analysis and composed the software code for data analysis. A. Parchure, and Y. Kim performed and analyzed secretion assays. O. Griesbeck provided Twitch sensors encoding plasmids. M. Schwartz provided the equipment. B. R. Ramazanov and J. von Blume wrote the paper.

## Abbreviations

ER: Endoplasmic reticulum
FRET: Fluorescence resonance energy transfer
nm: nanometer
TGN: Trans Golgi Network

## Supplementary data

### Supplementary Figure legends

**Supplementary 1. RUSH experiments using Cab45 clients demonstrate that delay in protein trafficking takes place at TGN**

(A) Representative histograms plotting fluorescence intensity distribution of COMP-GFP in cells at 30, 60, and 120 minutes (shown in red) during the RUSH experiment compared to timepoint “0” (shown in green overlay) in HeLa WT cells line treated with DMSO (upper row), HeLa WT cells treated with 2-APB (middle row), and HeLa VAPA/VAPB-dKO line (bottom row). 10.000 cells were analyzed for each condition. (B) Graph depicting mean normalized fluorescence intensity (F.I.) of RUSH LyzC-EGFP in BFP-GALNT1 (Golgi marker) region of interest (ROI) from and after the time when the maximal LyzC-EGFP F.I. is reached in the ROI (peak Golgi F.I.). The decrease in LyzC-EGFP F.I. represents the Golgi exit in the HeLa line treated with DMSO (black) versus 2-APB treated (red). At least 10 cells from independent experiments for each condition were analyzed with nonlinear regression (exponential function) indicated in the plot using GraphPad Prism9 built-in regression. (C) Graph depicting mean normalized fluorescence intensity (F.I.) of RUSH LyzC-EGFP in BFP-GALNT1 (Golgi marker) region of interest (ROI) from and after the time when the maximal LyzC-EGFP F.I. is reached in the ROI (peak Golgi F.I.). The decrease in LyzC-EGFP F.I. represents the Golgi exit in the HeLa line transfected with control siRNA (black) versus the Hela line transfected with siRNA to VAPA and VAPB (red). At least 10 cells from independent experiments for each condition were analyzed with nonlinear regression (exponential function) indicated in the plot using GraphPad Prism9 built-in regression.

**Supplementary 2. VAPA/VAPB dKO HeLa lines demonstrate delay in TGN export of Cab45 clients in RUSH experiments**

(A) Representative immunofluorescence images of RUSH experiments showing LyzC-GFP transport in HeLa WT and VAPA/B-dKO cell lines. Cells were transfected with KDEL-IRES-SBP-LyzC-EGFP and fixed at 0, 20, 40, and 60 min after adding biotin. Z-stack images (d = 0.2 μm) were analyzed. The arrowheads indicate cytoplasmic vesicles. Scale bars, 10 μm. (B) Representative immunofluorescence images of RUSH experiments showing COMP-GFP transport in HeLa WT and VAPA/B-dKO cell lines. Cells were transfected with KDEL-IRES-SBP-COMP-EGFP and fixed at 0, 20, 40, and 60 min after adding biotin. Z-stack images (d = 0.2 μm) were analyzed. The arrowheads indicate cytoplasmic vesicles. Scale bars, 10 μm. (C) The numbers of LyzC budding vesicles were quantified. The cytoplasmic vesicles were counted at each time point by analyzing z-stack images (d = 0.2 μm). The scatter dot plot represents the means ± SD of at least three independent experiments (n > 30 cells per condition). Statistical test, Kruskal–Wallis. (D) The numbers of COMP budding vesicles were quantified. The cytoplasmic vesicles were counted at each time point by analyzing z-stack images (d = 0.2 μm). The scatter dot plot represents the means ± SD of at least three independent experiments (n > 30 cells per condition). Statistical test, Kruskal–Wallis.

**Supplementary 3. MCS targeting FRET sensors localize with TGN markers.**

(A) Schematic representation of the domain structure of OSBP (top construct) and Twitch sensors fused proteins targeting ER-Golgi contact sites (N-PH-FFAT-Twitch and PH-FFAT-Twitch) as well as sensors targeting ER (fused with VAPA protein) and cytoplasmic version (two bottom constructs) used in this study. (B) IF images of HeLa lines stably expressing Twitch2b sensor targeting cytoplasm, ER and ER-TGN contact sites. (C) IF images of HeLa line expressing Twitch2b sensor targeting ER-TGN contact sites and its localization compared to TGN46 (TGN marker). (D) IF images of HeLa line expressing Twitch2b sensor targeting ER-TGN contact sites and its localization compared to GM130 (cis Golgi marker). (E) Dot plot representing FRET index for PH-FFAT-Twitch2b and PH-FFAT-Twitch7x, PH-FFAT-Twitch8x and PH-FFAT-Twitch9x within Golgi ROI of live cells and at steady state (***p<0.05); (F) Dot plot representing FRET index for PH-FFAT-Twitch2b and N-PH-FFAT-Twotch2b in cells at steady state. (G) Dot plot representing FRET index for N-PH-FFAT-Twitch2b cells at steady state and treated with 1uM ionomycin for 20 min (***p<0.05). (H) Dot plot representing FRET index for N-PH-FFAT-Twitch2b cells at steady state and treated with 1uM Histamine for 2 min(***p<0.05).

**Supplementary 4. Calibration procedures of Twitch2b sensor and fluorescence channels for FRET experiments**

(A) Schematic representation of the calibration experiment of Twich2b calcium sensor using Ca-EGTA buffers in live cells (Please see Materials and Methods section for detailed description). (B) Pseudocolor heatmap of FRET index for CYTO-Twitch2b during calibration experiment. Pseudocolor bar FRET index value in range 0-0.4. (C) Calibration of fluorescence channels for Twitch sensor using HeLa lines expressing PH-FFAT-mCerulean3 (only FRET donor), HeLa lines expressing PH-FFAT-cpVenus (only FRET acceptor)) and PH-FFAT-Twitch2b (FRET pair with calcium binding domain) sensor.

## Table legends

**Supplementary Table1.** Twitch sensors and their affinity to Ca^2+^ ions.

| FRET Sensor | Calcium affinity of TnC (KD) |
| --- | --- |
| Twitch 2b | 200 nM |
| Twitch 7x | 51.5 uM |
| Twitch 8x | 139 uM |
| Twitch 9x | 174 uM |

**Supplementary Table2.**
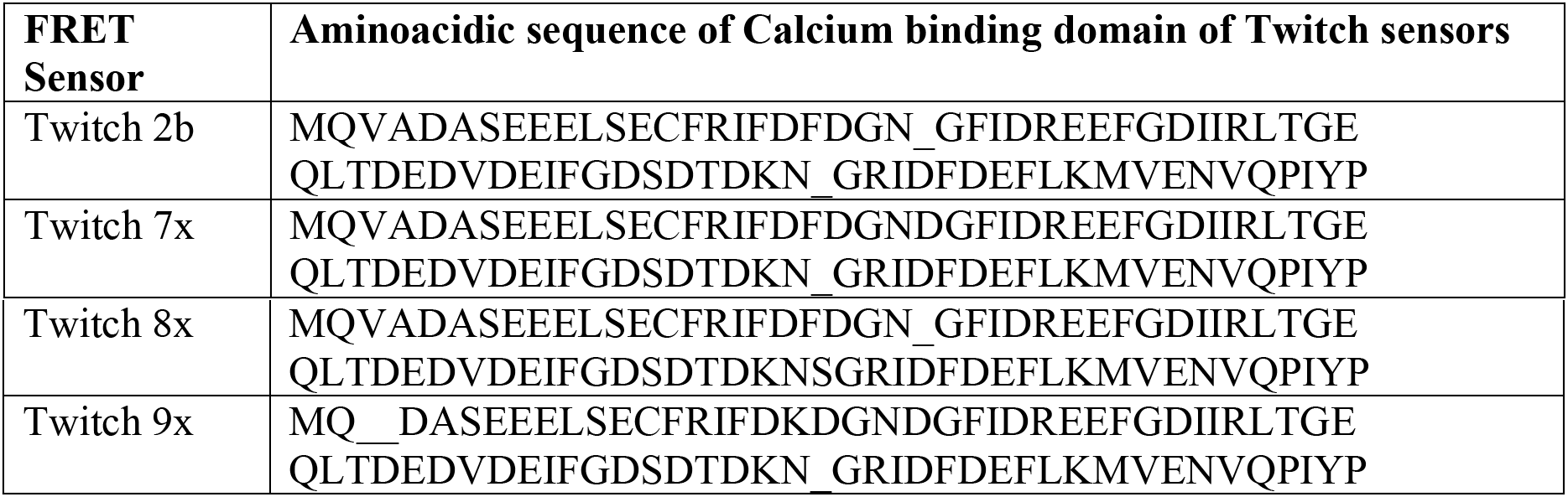
Twitch sensors and aminoacidic sequences of calcium-binding domain.

