## Supplementary figures and images for "Calcium flux through ER-TGN contact sites facilitates cargo export"

### Supplementary figure 1

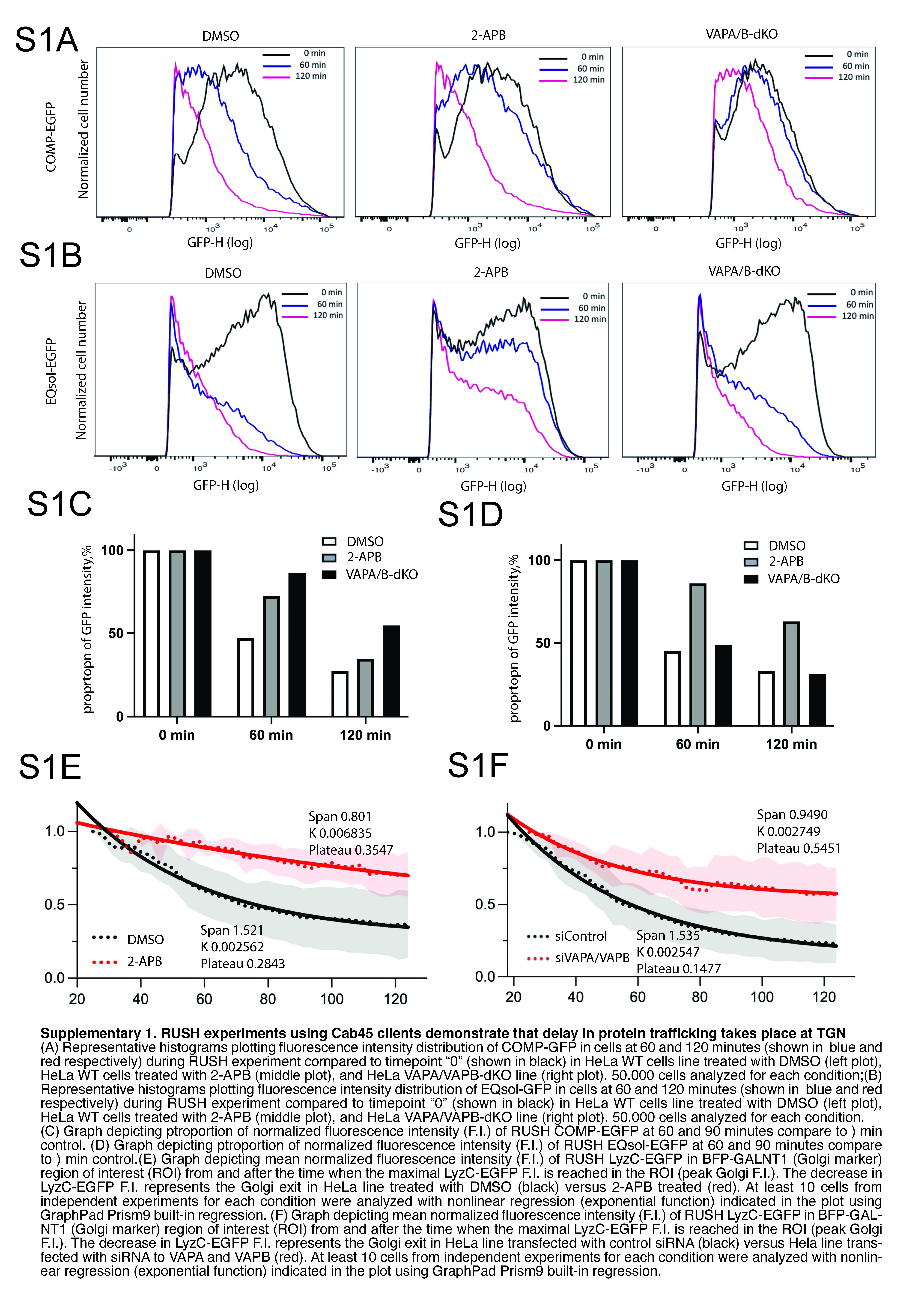

### Supplementary figure 2

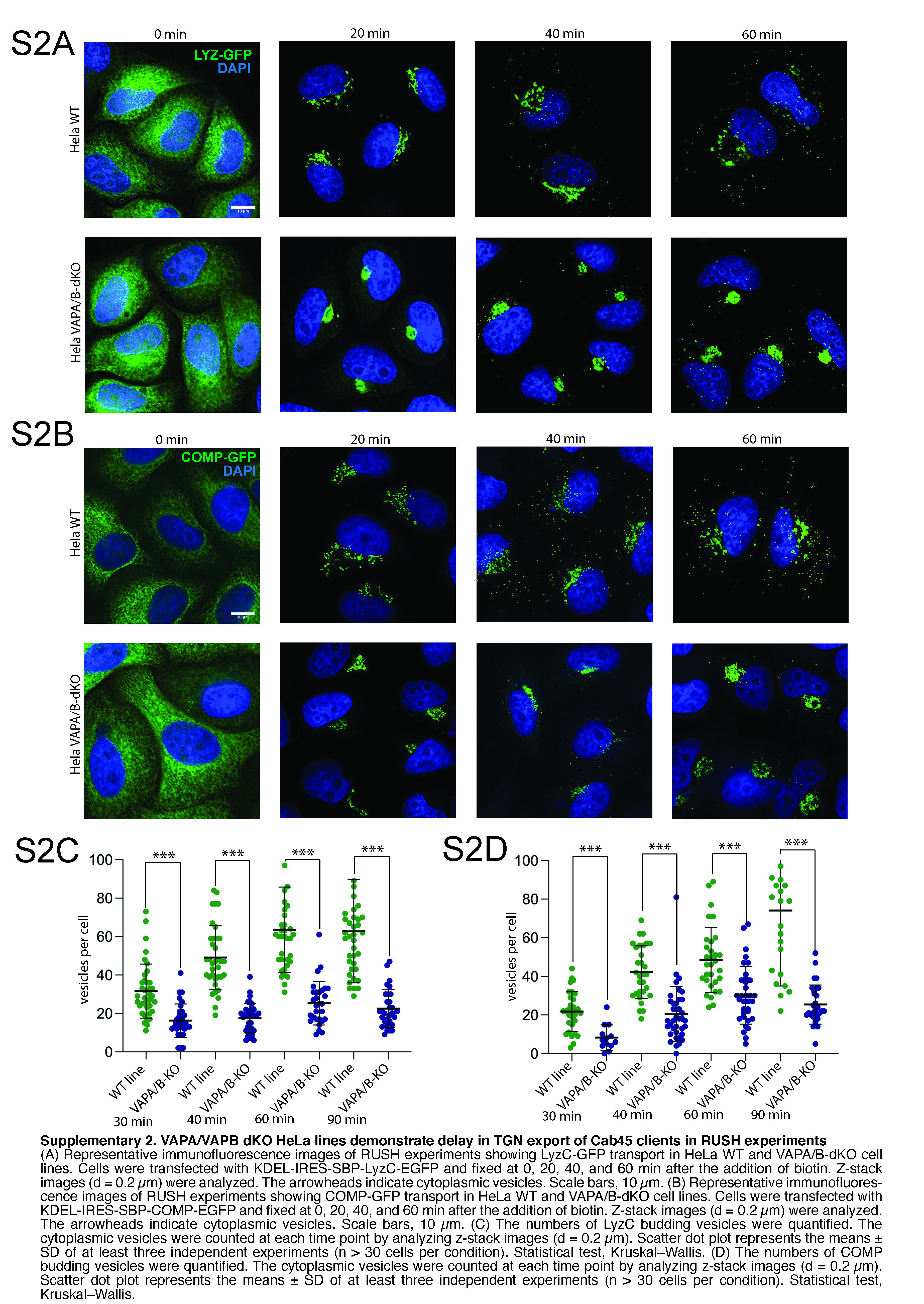

### Supplementary figure 3

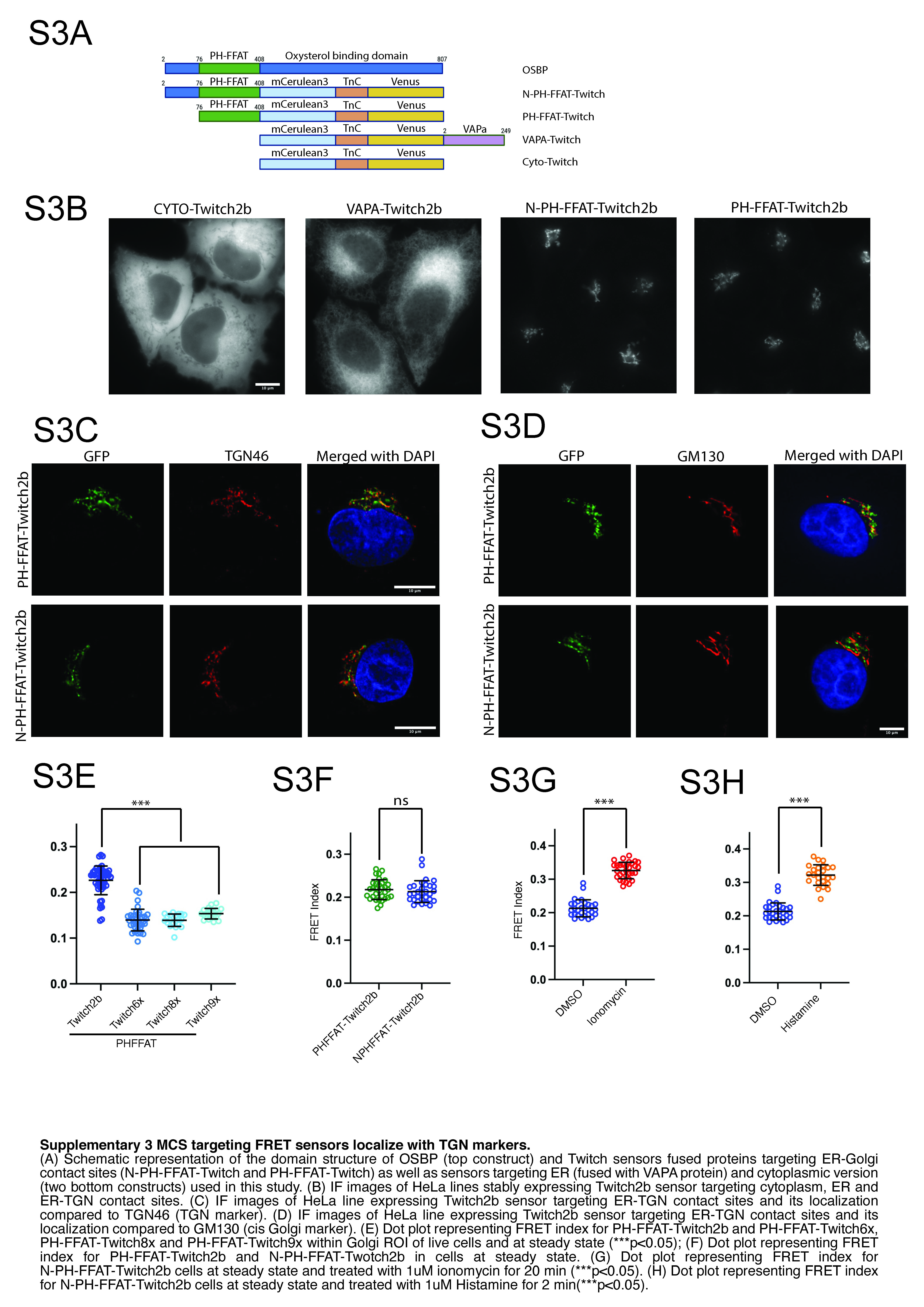

### Supplementary figure 4

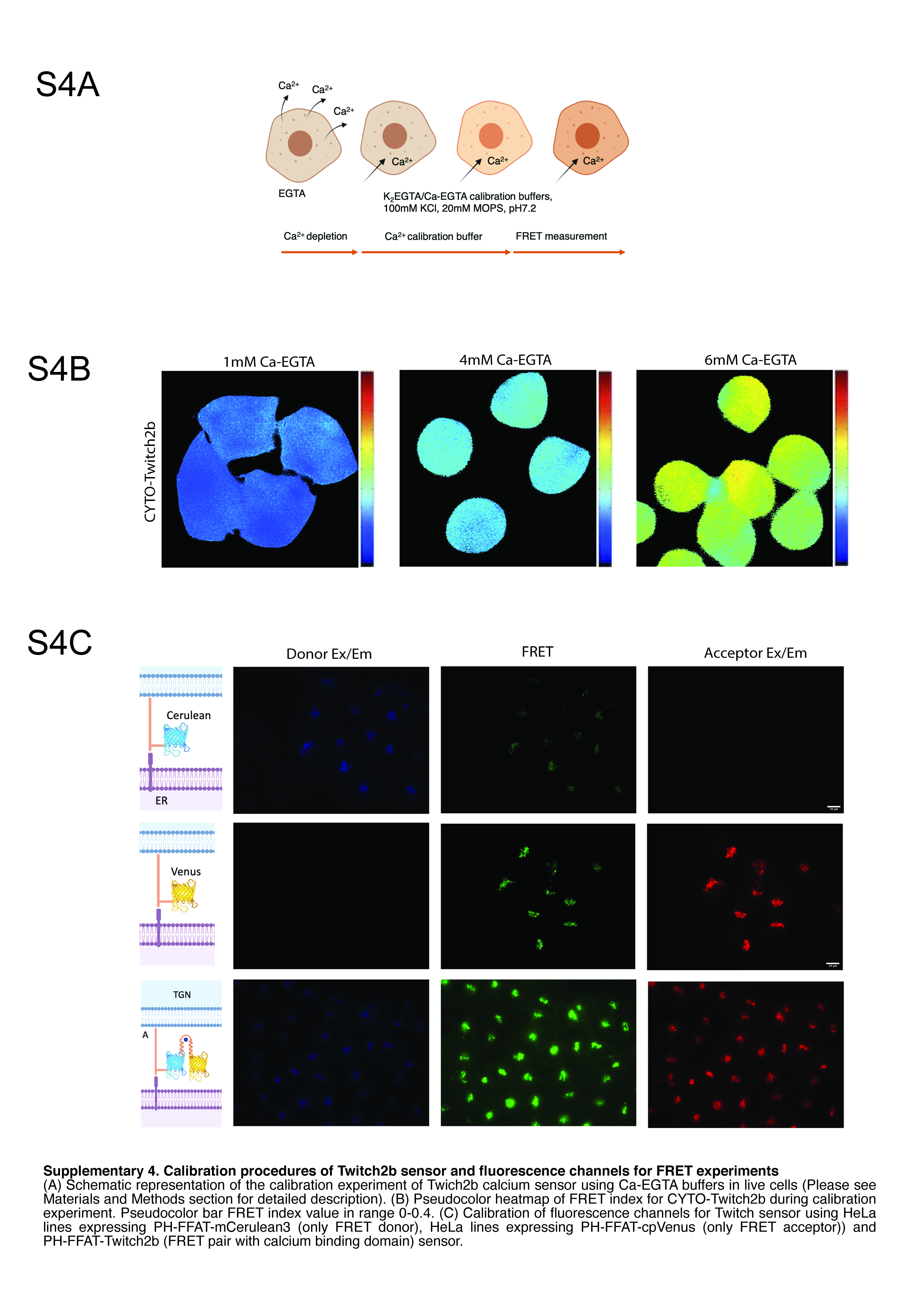
